# DNA sequencing with stacked nanopores and exonuclease: a simulation-based analysis

**DOI:** 10.1101/038034

**Authors:** G. Sampath

## Abstract

Experiments (Clarke et al., *Nat. Nanotech*., 2009, **4**, 265-270) have shown that DNA could be sequenced using a nanopore-based electrolytic cell in which an exonuclease enzyme in the *cis* chamber cleaves the leading base of a strand of DNA. The base is identified (with a reported accuracy that exceeds 99%) by the level of the current blockade it causes in the pore; a biological adapter inside slows down the base to lower the detection bandwidth required. This approach, which has been mathematically modeled, analyzed, and simulated (Reiner et al., *J. Chem. Phys.,* 2012, **137**, 214903; Brady and Reiner, *ibid.,* 2015, **143**, 074904), is error-prone because bases may be lost to diffusion or enter the pore out of order. Here a modified cell with three stacked nanopores (UNP, MNP, and DNP) and the enzyme attached to the *trans* side of UNP is proposed. Translocation of a base is simulated with the random walk of a dimensionless particle; the results show that bases translocate through MNP and DNP in sequence order without loss. If this holds in practice then with a suitably designed adapter and compatible enzyme turnover rates base calling accuracy would be limited only by the accuracy of base discrimination. Potential implementation issues are discussed.

## 1. Exonuclease sequencing of DNA

In contrast with most other next generation methods of DNA sequencing, nanopore-based methods require a minimal amount of preparation and do not use labeling of any kind. Sequencing is based on an electrolytic cell with the structure [*cis,* membrane-with-nanopore, *trans*] and a potential difference between the *cis* and *trans* chambers. Negatively charged bases (A, T, C, G) in a strand of DNA are identified by the level of current blockade that results when a base passes through the pore. Pores may be biological (αHL, MspA) or synthetic (silicon nitride, silicon oxide, graphene, molybdenum sulphide, etc.). See [1,2] for recent reviews.

In ‘strand sequencing’ [3] base identification is based on an intact strand of DNA passing from *cis* to *trans.* Discrimination among the bases is limited by pore length, speed of translocation of the strand, and efficiency of deconvolution (that is, extraction of signal levels for individual bases from the measured signal). In a variant known as ‘sequencing by synthesis’ (SBS) a polymerase on the *cis* side is used to slow down the strand [4].

In ‘exonuclease sequencing’ [5] an exonuclease enzyme covalently bonded to the pore on the *cis* side successively cleaves the leading base (actually ‘mononucleotide’; it is common, although not accurate, practice to use the two terms interchangeably in this context) from the strand. Blockade signals for individual bases being separated in time and space no signal deconvolution is necessary, so base calling, especially with homopolymers (sequences of identical bases), is, ideally, not the problem it is in strand sequencing. A covalently attached cyclodextrin adapter in the pore slows down the base and reduces the detector bandwidth required. Bases are identified with high accuracy (over 99%, as reported in [5]). However, the process is error-prone because a cleaved base may diffuse back into the *cis* bulk and be missed or called out of order [6].

To remedy this a tandem cell [7] with the structure [cis1, upstream pore (UNP), *trans1/cis2,* downstream pore (DNP), *transi]* and the exonuclease enzyme covalently bonded to the downstream side of UNP was proposed recently. A Fokker-Planck model was used to show that cleaved bases are released in *trans1/cis2* without regression into cis1 and, under the right conditions, reach DNP in order. However due to the large μm) size of *trans1/cis2* the possibility exists of diffusion losses in *trans1/cis*2. This report proposes a modified version that is aimed at resolving this problem.

## 2. The present work

The μm-sized *trans1/cis2* chamber in a tandem cell is replaced with a membrane 30-50 nm thick containing an hourglass-shaped nanopore. Such nanopores have been fabricated in the laboratory with a synthetic material such as silicon nitride or silicon oxide [1,2]. The cell now has the structure [cis, upstream pore (UNP), middle pore (MNP), downstream pore (DNP), *trans*]. See Figure 1.

**Figure 1.**
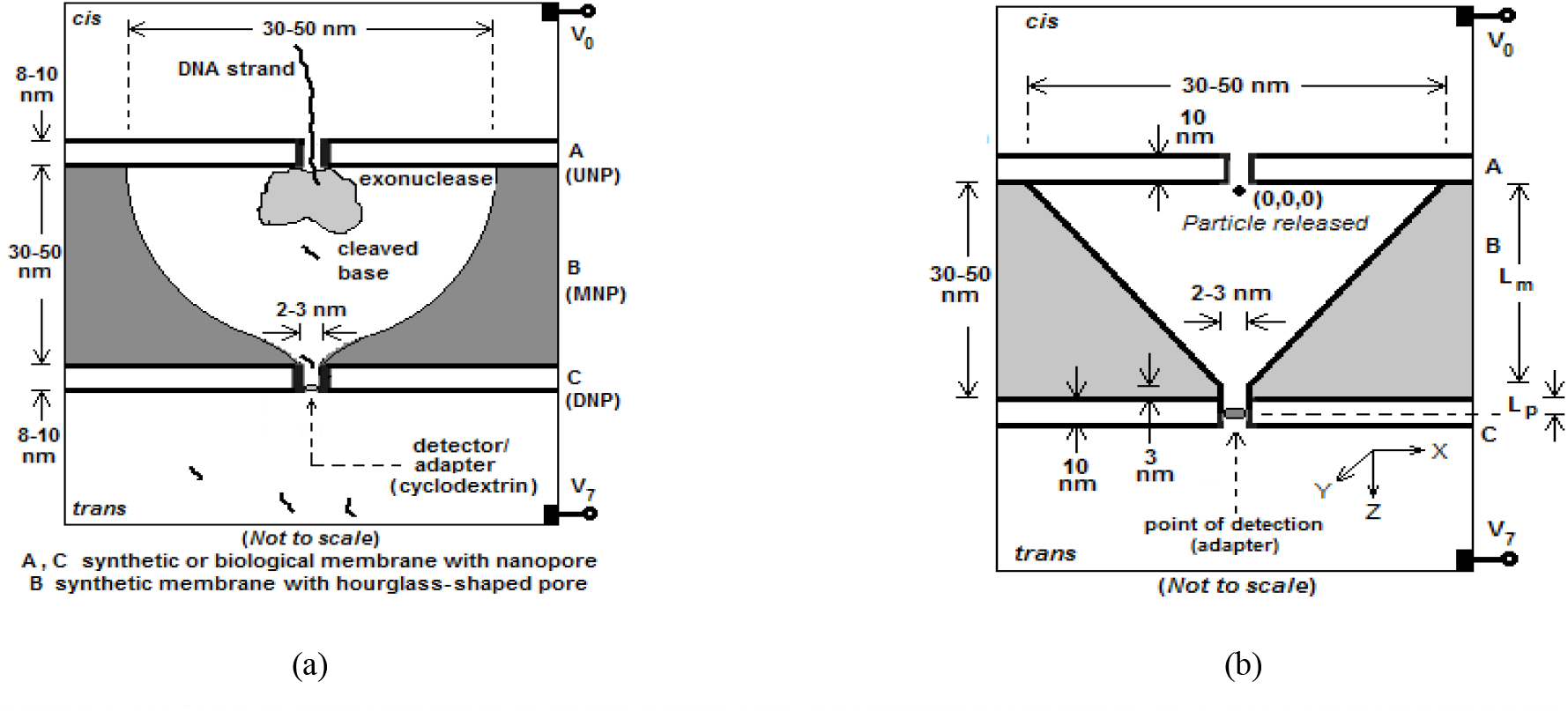
(a) Stack of three nanopores in an electrolytic cell; (b) Geometry (symmetric in X and Y) used in simulation UNP: upstream pore, MNP: middle pore, DNP: downstream pore. Adapter covalently attached near bottom of DNP.

The Fokker-Planck model developed earlier [7] for the tandem cell can be applied individually to DNP and MNP in the proposed structure. It is a good representation of the diffusion-drift process inside DNP, while inside MNP it is a reasonable approximation. Simulation studies, detailed in Section 5 below, indicate that there is no loss of cleaved bases to diffusion in MNP, and that cleaved bases reach and translocate through DNP in sequence order by a combination of diffusion and drift. If a suitably designed adapter and matching exonuclease turnover rates are used, sequencing accuracy would be determined only by base discrimination capability, which was shown in [5] to have a better than 99% accuracy. (This assumes a Gaussian fit of the data. If a Voigt distribution is used instead the accuracy drops to 92% [6], which is still high when compared with the measured accuracy of most strand sequencing methods.) The proposed structure may be fabricated using techniques currently available for biological and synthetic pores and assembled by stacking individually fabricated pores. Potential implementation issues are discussed in Section 7.

## 3. Model of cell with three stacked nanopores and exonuclease

In modeling the proposed structure the following assumptions are made:

1. The electric field near the entrance to the first pore is high enough that a strand of DNA (uncoiled and without any secondary structure) is pulled into UNP [8];
2. A base cleaved by the enzyme on the *trans* side of UNP cannot regress into UNP (and therefore *cis)* because it is blocked by the remaining DNA strand in UNP;
3. For simplicity the pore at the bottom of MNP is merged with DNP and the result considered to form a single pore, with the rest of the hourglass constituting MNP;
4. With a high enough potential difference across DNP, a cleaved base after encountering the adapter in DNP always translocates into *trans* [5] and does not permanently regress into DNP and thence into MNP. Thus while there may be some oscillation across the interface between DNP and *trans* the net movement of the particle is ultimately always into *trans*;
5. A cleaved base is a dimensionless particle carrying a negative electric charge; in MNP it is subject only to diffusion and drift due to electrophoresis, while in DNP it is subject to diffusion and drift due to electrophoresis as well as electro-osmotic effects from interaction with an ion-selective pore [6];
6. The walls of MNP and DNP are reflective, and the enzyme itself is assumed to have no effect on the trajectory of a cleaved base;
7. Hydrodynamic effects in both MNP and DNP are neglected;
8. Exonuclease cleavage efficiency in MNP is the same as in the bulk (for a study of Exonuclease I in the bulk see [9]).

Based on the above assumptions the Fokker-Planck model described in [7], which incorporates diffusion and drift (due to electrophoresis and electro-osmotic effects), can be adapted to the structure in Figure 1a. (This model is similar to one based on the original exonuclease sequencing approach [6] and recently extended in [10].) However the structure cannot be solved analytically because of the discontinuity between MNP and DNP; solution requires matching of boundary conditions between the two domains, which cannot be done analytically. There are two alternatives available to deal with the discontinuity: numerical solution and simulation. In the present work the latter is used.

## 4. Conditions for accurate sequencing

With the proposed structure, accurate sequencing of a strand of DNA requires the following conditions to be satisfied:

C1: Cleaved bases must enter DNP from MNP in sequence order;
C2: No more than one cleaved base may occupy DNP at any time;
C3: The detector circuit has a sufficiently high bandwidth that blockade events are not missed.

Let the interval between successive cleavages by the exonuclease be T_exo_, and the corresponding minimum and mean T_exo-mm_ and T_exo-mean_. For a cleaved base let T_MNF_ = translocation time in MNP, T_DNP_ = translocation time in DNP, and T_adapter_ = translocation delay in the adapter. Let the corresponding minimum, maximum, and mean be T_MNP-mm_, T_MNP-max_, T_MNP-mean_, T_DNP-min_ T_DNP-max_, T_DNP-mean_ T_adapter-min_, T_adapter-max_, and T_adapter-mean_. Let T_detectorr_ be the pulse width resolution of the detector.

C_1_ is strongly satisfied if

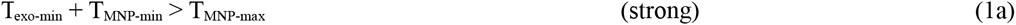

It is weakly satisfied if

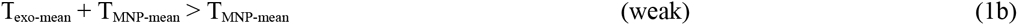

Similarly, for C_2_:

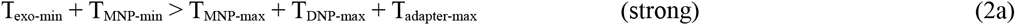

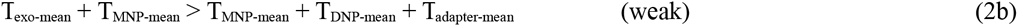

On comparison with Equations 1a and 1b, C_2_ is seen to subsume C_1_.

For C_3_ to be satisfied, the detector’s pulse resolution time must be smaller than the time taken by a base to translocate through DNP and the adapter into *trans.* This requires

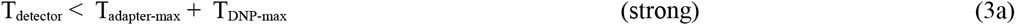

or, in a weaker version,

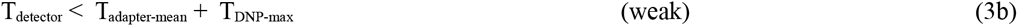

Conversely, the maximum acceptable delay due to the adapter is given by

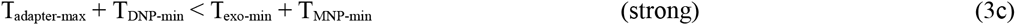

On the other hand a minimum limit is placed on adapter delays by the bandwidth (or, equivalently, pulse width resolution) of the detector circuit. Presently the available bandwidth is about 2 MHz or time resolution of ∽1 μs [11]. (Thus with the fastest detectors available at present adapter delays have to be at least 1 μs long.) Equation 3c then becomes

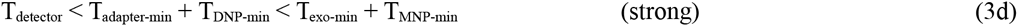

In principle C2 can be relaxed. Thus successive cleaved bases may enter DNP and queue up at the adapter to be recognized in succession as the adapter holds and releases them on the *trans* side in sequence order. Exosequencing then reduces to strand sequencing. In this case, while heteropolymers will be recognized correctly because of the differences in their blockade levels, the advantage that exosequencing has over strand sequencing in recognizing homopolymers from their identical blockade levels due to the time separation present in the absence of such telescoping at the adapter is lost.

## 5. Simulation: assumptions, measures of interest, calculations, choice of parameters

The Fokker-Planck equation incorporates two kinds of terms, one due to diffusion and the other due to drift [7]. The former corresponds to random movements in 3-d space, while the latter is movement in a directed electric field that is assumed to be constant in the region simulated. An additional force exists in a narrow region like DNP due to electroosmotic effects inside an ion-sensitive pore; this can be approximated by an equivalent drift field coaxial with DNP. The velocities due to the two drifts are added. Such diffusion and drift of particles in confined regions like MNP and DNP is easily simulated.

The proposed structure was simulated to determine how well the conditions in Section 4 are satisfied. Simulation runs were designed to cover a range of parameters, including pore dimensions, diffusion constant, mobility, drift velocity due to electrophoresis and electro-osmotic effects, and pore voltage. Particle trajectories were simulated in the continuum defined by MNP and DNP (Figure 1b) using standard procedures, and various performance measures for use in assessing the proposed structure computed. The simulation protocol used closely follows the one described in [6] and modified in [10]. (In the latter, trajectories of cleaved bases in the original exosequencing method using a conventional electrolytic cell with a single pore were simulated using diffusion constants that are different in the pore and in *cis.* This has been done in the present work as well. Variations in the protocol are noted where appropriate.)

*Assumptions and simplifications*

1. The hourglass shape of MNP is approximated by a tapered cylinder ending in a nanopore that is merged with DNP as shown in Figure 1b.
2. The potential difference of V_cia-trans_ applied between Ag/AgCl electrodes situated at the top of *cis* and bottom of *trans* divides as V_cis_ over *cis,* V_UNP_ across UNP, V_MNP_ across MNP, V_DNP_ across DNP, and V_trans_ over *trans*; potential differences across MNP and DNP are assigned approximate values using the assumed conductances of the two pores.
3. A cleaved base is a dimensionless particle that is reflected off the walls of MNP and DNP.
4. In MNP the particle experiences random diffusive movement and directed drift due to the electric field E_MNP_ (resulting from the applied potential V_MNP_) in the 3-d space defined by MNP; in DNP it experiences the electric field E_DNP_ that exists over the 1-d element representing DNP as well as electro-osmotic effects due to interaction of the base with the walls of an ion-sensitive pore, the related calculations are given below.
5. For the voltages assumed, the equivalent electro-osmotic potential is considerably smaller in comparison. For simplicity therefore, the field due to an electro-osmotic field is subsumed under E_DNP_ in the simulation.
6. The diffusion constant is smaller in DNP than in MNP because the environment in DNP is ‘crowded’. The reverse case also has been simulated, similar to [10].
7. The particle is considered detected (‘trapped’ and released on the *trans* side) when it reaches the adapter, which is positioned at or near the bottom of DNP and plays the role of a detector [5]. The potential difference V_DNP_ is sufficient for the ‘trapped’ particle to be released and then translocate into *trans* without regressing into DNP [5].

*Measures of interest*

The following measures are of interest:

1. Minimum, maximum, and mean translocation time through MNP;
2. Minimum, maximum, and mean translocation time through DNP;
3. Number of particles ‘lost’.

(A particle is considered lost if the number of random walk steps exceeds 2.4 × 10^9^, which, with a time step of 10^−12^ sec corresponds to a total trajectory time of 2.4 ms.)

*Constants used*

1. Diffusion constant 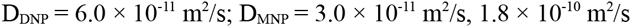
2. Mobility constant 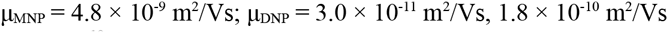
3. Electron charge e 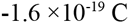

*Calculations*

1. The electric field across a pore of interest is E = V/L, where V is the applied voltage across the pore and L is the thickness (height) of the pore. The electrophoretic drift velocity is v_epd_ = μΕ where μ is the mobility and E the electric field.
2. Following [6] the equivalent electro-osmotic drift velocity v_eo_ of a charged base in an ion-sensitive DNP is given by

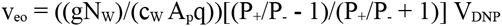

where g = nanopore conductance = 1 nS, N_w_ = number of water molecules hydrating each ion = 10, c_w_ = molar concentration of water = 3.4 × 10^28^ m^-3^, A_p_ = average cross-sectional area of pore = 6 × 10^−18^ m^3^, q = charge on ion = 2e, P+/ P-= nanopore ion selectivity ratio = 0.1, and V_DNP_ = applied potential difference across DNP. V_eo_ being small compared to the electrophoretic drift velocity V_DNP_, it is subsumed into vepd = V_DNP_ in the simulations for simplicity.
3. The displacement of a particle inside a pore is the vector sum of the displacements due to diffusion, electrophoresis, and electro-osmosis. Diffusive movement occurs in a random direction in 3-d space and is simulated as a displacement during a small time interval Δt (set to the simulation time step of 1 picosecond = 10^−12^ s). The diffusion displacement size is √(6DΔt), where D is the diffusion constant of the medium. A random direction is generated as a unit vector on the surface of a sphere. This vector is then multiplied by the step size and the translation due to drift in the direction of the electric field (E_MNP_ or E_DNP_) is added to it. The resulting position is accepted if it is inside MNP or DNP and rejected otherwise.

*Simulation procedure*

The following procedure is repeated N times with a time step of 10^−12^ s.

**Procedure** Simulate-Particle-Trajectory

1. The particle is released at **p** = (x,y,z) = (0,0,0); see Figure 1b for coordinate system used. Initialize the number of simulation steps and the number of particles lost to diffusion both to 0.
2. Generate a random vector r to the surface of the unit sphere centered at the current position. Multiply this by the diffusion step size, which is determined by the current position. If **p** is in MNP the size is √(6D_MNP_ Δt), if **p** is in DNP the size is √(6D_DNP_ Δt), with Δt = 10^−12^ s.
3. Calculate the drift velocity V_MNP_ or V_DNP_ depending on whether **p** is in MNP or DNP, and thence the drift displacement in the z direction as **d** = (0, 0, V_MNP_ Δt) or (0, 0, V_DNP_ Δt).
4. If **p** + **r** + **d** is in MNP or DNP set **p** to **p** + **r** + **d** and increment the number of simulations by 1; else reject the move. (The latter flows from the assumption that the walls of MNP and DNP are reflective. The decision to reject is not quite accurate, but as the number of reflections at the wall << the number of steps not reflected the error can be neglected.)
5. If the particle has reached the bottom of DNP, it has been detected so exit.
6. Add 1 to the number of simulation steps; if the result is greater than the prescribed maximum number of steps (the stopping criterion for the simulation, here set to 2.4 × 10^9^) add 1 to the number of particles lost to diffusion and exit.
7. Go to Step 2.

*Choice of parameters in simulations*

1. For MNP two different sizes were used: thickness 40 nm, width 40 nm; thickness 30 nm, width 30 nm. These are in the range of sizes of silicon nitride membranes [1,2] with hourglass-shaped nanopores. For DNP two thicknesses of 10 nm and 8 nm, which are in the range of biological nanopores (α-hemolysin and MspA) [1,2], were used. A pore width of 2.4-3.0 nm was used, consistent with the dimensions of α-hemolysin and MspA.
2. Voltages of 150 mV and 180 mV across DNP were used, these are within the range of most nanopore sequencing studies. Lower voltages are not used as they could result in regression of a detected base into DNP from the adapter [5]. (Also it would result in lower ionic currents, which could adversely affect the detection of blockades.) In a conventional electrolytic cell with a nanopore, the drop across *cis* and *trans* is less than 1-2% of the voltage across DNP [6], and many models and simulation-based studies, for example [6] and [10], assume it to be zero. To allow for both possibilities the voltage across MNP was assumed to be 1% of the voltage across DNP in three of the five runs (with the resulting electric field taken to be constant and uniform over MNP) and 0 in the remaining two runs.
3. Based on the bulk diffusion constant of KCl D_bulk_ = 3.0 × 10^−10^ m^2^/s, D_DNP_ was set to 0.2 D_bulk_ (to simulate the ‘crowded’ environment in DNP [6]). For MNP two values were used: 3 D_DNP_ and 0.5 D_DNP_. The last value was used to simulate the effect of using a gel to slow down a translocating base [12,13] (or, equivalently, the use of alternative electrolytes [14], with a potential slowdown factor of 7 to 11).

## 6. Simulation results and analysis

Each run consisted of executing the procedure in Section 5 N = 1000 times for five different sets of parameters. The statistics generated are summarized in Table 1.

**Table 1.**
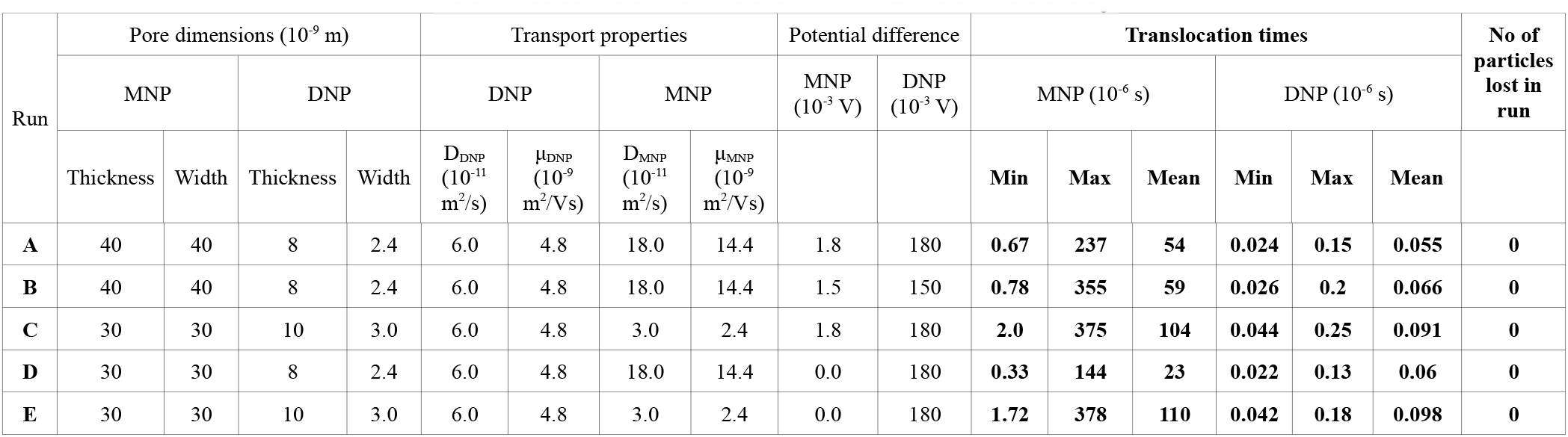
Results of simulations: statistics summary

Data from [9], [5] and [11] have been used in evaluating the above results:

1. Exonuclease turnover intervals between two cleavages [9]:

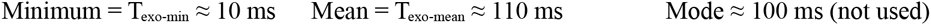 These values are for the enzyme cleaving bases from a DNA strand in the bulk electrolyte. In the absence of data these values are assumed to be valid inside MNP.
2. Mean dwell times (τ_off_) for bases in the adapter [5]:

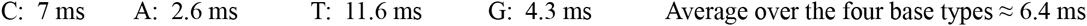 In the calculations below the average value of 6.4 ms is used for T adapter-mean (= Toff).
3. Detector resolution time [11]:

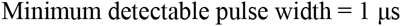 Conditions C_1_ through C_3_ are now checked using the above data and the smallest minimum and largest maximum over columns 12 through 17 in Table 1.

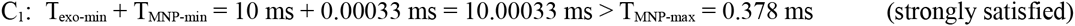

Two successive cleaved bases enter DNP in sequence order.

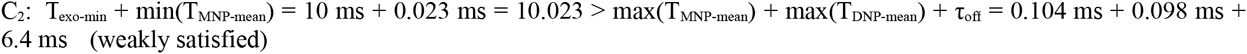

Thus on the average no more than one base occupies DNP at any time.

C_3_: Since only mean values are available for T_adapter_ a weak form of C_3_ is checked:

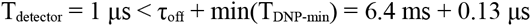

and

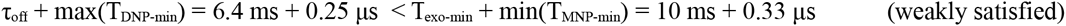

Thus on the average blockade events are not missed.

On the other hand, if the mean adapter delay of 11.6 ms for thymine (T) [5] is considered, C3 is not satisfied, so the adapter would have to be designed to have a smaller delay.

In passing it may be noted that the exonuclease cleavage interval (turnover rate) itself can be controlled by controlling the electrolyte pH, temperature, and related factors [9]).

## 7. Notes on potential implementation

Several recent reports describe systems with two nanopores in tandem. One of these is used to measure mobility and particle sizes to identify specific proteins, but the nanopores are comparatively larger, with dimensions in the 10’s of nanometers [15]. A commercial website [16] mentions the use of two pores to control translocation of DNA with a time-varying potential difference, but no details are available. Another report [17] describes the use of two stacked pores to measure the free-solution mobility of DNA molecules based on the time taken to translocate from one pore to the other. In one recent study two pores were brought together to form a ‘cage’ to trap DNA and cut it into fragments using a restriction endonuclease [18]. The number of fragments and their sizes were obtained from blockade pulses created by applying a reverse voltage. Fabrication of stacks of single-atom-thick graphene layers has also been reported [19].

Figure 2 shows possible implementations of the scheme proposed in this report. Figure 2a is based on a hybrid biological-synthetic-biological structure. A variation of this consists of flipping the first nanopore vertically so that the broader vestibule of the oHL pore provides a platform to attach the exonuclease enzyme. In Figure 2b an oHL pore is inserted into a pore in a synthetic membrane (see next paragraph). The oHL in membrane A has an exonuclease enzyme covalently attached or fused to it.

**Figure 1.**
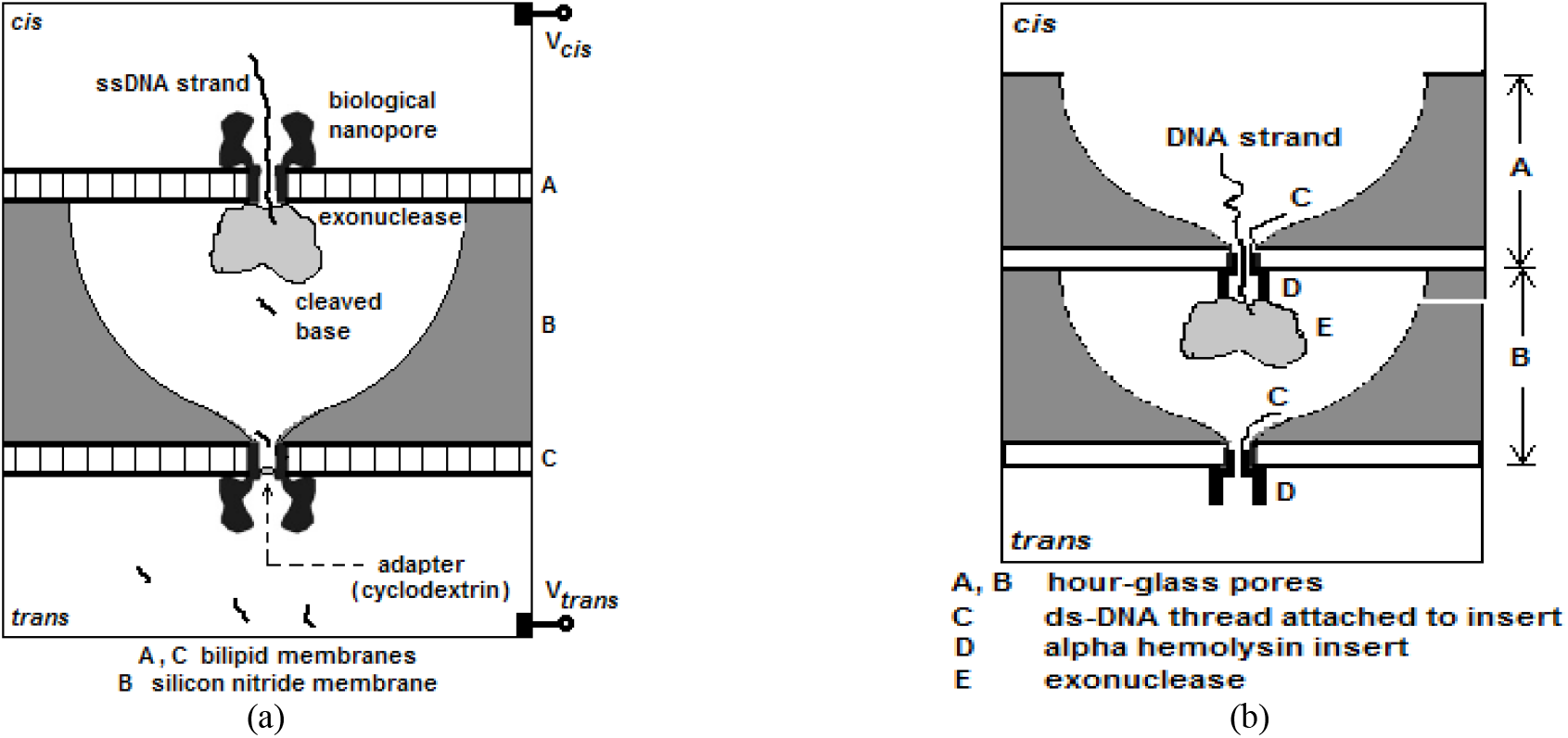
(a) Implementing a stack of three nanopores using a biological-synthetic-biological pore assembly (b) Structure with two synthetic membranes with hourglass-shaped pores and oHL pores inserted into them

Biological pores have the disadvantage that they are embedded in bilipid layers that are mechanically unstable. An alternative approach dispenses with the bilipid layers and uses a synthetic pore to support an oHL pore. The latter is inserted into a silicon nitride membrane with an hourglass-shaped pore [20]. The report mentions a success rate of 20-40% for the insertion but does not say if the insertion is dependably permanent. This method may be useful for the scheme proposed here if the procedure is reliable. It also requires the exonuclease to be covalently attached or fused to the alpha hemolysin before insertion. With such an approach the structure presented here can be implemented as two hybrid pores A and B stacked one on top of the other. Such stacking can be done by mechanical assembly in which MNP is aligned with UNP; the alignment is made easier by the hourglass shape, whose broad mouth can also accommodate exonuclease molecules that are covalently attached to the *trans* side of pore A.

## References

[1] M. Wanunu, “Nanopores: a journey towards DNA sequencing.” Phys Life Rev, 2012, 9, 125–158.

[2] Y. Wang, Q. Yang, and Z. Wang, ‘The evolution of nanopore sequencing,’ Front. Genet., 2015, 5, 449.

[3] J. J. Kasianowicz, E. Brandin, D. Branton, and D. W. Deamer, "Characterization of individual polynucleotide molecules using a membrane channel," PNAS, 1996, 93, 13770–13773.

[4] K. R. Lieberman, G. M. Cherf, M. J. Doody, F. Olasagasti, Y Kolodji, and M. Akeson, "Processive replication of single DNA molecules in a nanopore catalyzed by phi29 DNA polymerase," J. Am. Chem. Soc., 2010, 132, 17961–72.

[5] J. Clarke, H-C. Wu, L. Jayasinghe, A. Patel, S. Reid, and H. Bayley, "Continuous base identification for single-molecule nanopore DNA sequencing", Nature Nanotech., 2009, 4, pp. 265–270.

[6] J. E. Reiner, A. Balijepalli, J. F. Robertson, D. L. Burden, B. S. Drown, and J. J. Kasianowicz, “The effects of diffusion on an exonuclease/nanopore-based DNA sequencing engine”, J. Chem Phys. 2012, 137, 214903.

[7] G. Sampath, “A tandem cell for nanopore-based DNA sequencing with exonuclease,” RSC Adv., 2015, 5, 167–171.

[8] A. Y Grosberg, Y Rabin. “DNA capture into a nanopore: interplay of diffusion and electrohydrodynamics,” J. Chem. Phys., 2010, 133, 165102.

[9] J. H. Werner, H. Cai, R. A. Keller, and P M. Goodwin, “Exonuclease I hydrolyzes DNA with a distribution of rates”, Biophys. J., 2005, 88, pp. 1403–1412.

[10] K. T. Brady and J. E. Reiner, “Improving the prospects of cleavage-based nanopore sequencing engines,” J. Chem. Phys., 2015, 143, 074904.

[11] J. K. Rosenstein, M. Wanunu, C. A. Merchant, M, Drndic, and K. L. Shepard, ‘Integrated nanopore sensing platform with submicrosecond temporal resolution,’ Nat. Methods, 2012, 9, pp 487–492.

[12] Z. Tang, Z. Liang, Bo Lu, Ji Li, Rui Hu, Qing Zhao, and Dapeng Yu, “Gel mesh as ‘brake’ to slow down DNA translocation through solid-state nanopores,” Nanoscale, 2015, 7, 13207–13214.

[13] M. Waugh, A. Carlsen, D. Sean, G. W. Slater, K. Briggs, H. Kwok, and V Tabard-Cossa, “Interfacing solid-state nanopores with gel media to slow DNA translocations,” Electrophoresis, 2015, 36, 1759–1767.

[14] C. Plesa, N. van Loo, and C. Dekker, “DNA nanopore translocation in glutamate solutions,” Nanoscale, 2015, 7, 13605.

[15] Z. D. Harms, D. G. Haywood, A. R. Kneller, L. Selzer, A. Zlotnick, and S. C. Jacobson, "Single-particle electrophoresis in nanochannels," Anal. Chem., 2014, DOI: 10.1021/ac503527d.

[16] http://twoporeguys.com/technology.html. (Accessed January 15, 2016)

[17] M. Langecker, D. Pedone, F. C. Simmel, and U. Rant, "Electrophoretic time-of-flight measurements of single DNA molecules with two stacked nanopores," Nano Lett., 2011, 11, 5002–5007.

[18] X. Liu, M. M. Skanata, and D. Stein, “Entropic cages for trapping DNA near a nanopore,” Nat. Communs. 2015, doi:10.1038/ncomms7222.

[19] S. Garaj, S. Liu, J. A. Golovchenko, and D. Branton, “Molecule-hugging graphene nanopores,” PNAS, 2013, 110, 12192–6.

[20] A. R. Hall, A. Scott, D. Rotem, K. K. Mehta, H. Bayley, and C. Dekker, “Hybrid pore formation by directed insertion of a haemolysin into solid-state nanopores,” Nature Nanotech., 28 November 2010, DOI: 10.1038/nnano.2010.237.

